# Distinct and stage-specific contributions of TET1 and TET2 to stepwise cytosine oxidation in the transition from naive to primed pluripotency

**DOI:** 10.1101/281519

**Authors:** Christopher B. Mulholland, Franziska R. Traube, Edris Parsa, Eva-Maria Eckl, Maximillian Schönung, Miha Modic, Michael D. Bartoschek, Paul Stolz, Joel Ryan, Thomas Carell, Heinrich Leonhardt, Sebastian Bultmann

## Abstract

The TET-oxidized cytosine derivatives, 5-hydroxymethylcytosine (5hmC) and 5-formylcytosine (5fC), are considered DNA demethylation intermediates as well as stable epigenetic marks in mammals. We compared modified cytosine and enzyme levels in TET-knockout cells during naive pluripotency exit and found distinct and differentiation-dependent contributions of TET1 and TET2 to 5hmC and 5fC formation. The divergent modified cytosine levels argue for independent consecutive oxidation steps *in vivo* with broad implications for epigenetic regulation.

## MAIN TEXT

DNA methylation is required for epigenetic regulation of gene expression and genome stability in mammals (Smith and Meissner, 2013). Recently, methylated cytosine (5mC) was discovered to be oxidized by the Ten-eleven Translocation (TET) family of dioxygenases (Tahiliani et al., 2009). The three mammalian homologs, TET1, TET2, and TET3, share a conserved dioxygenase domain and catalyze the stepwise oxidation from 5mC to 5-hydroxymethylcytosine (5hmC), 5-formylcytosine (5fC), and 5-carboxylcytosine (5caC) (Figure 1a) (He et al., 2011; Ito et al., 2011; Iyer et al., 2009; Pfaffeneder et al., 2011). These oxidized cytosine derivatives have been shown to be intermediates of passive and active DNA demethylation (He et al., 2011; Ito et al., 2011), yet may also serve as stable epigenetic marks (Bachman et al., 2014, 2015). Moreover, their largely separate genomic distributions and reader proteins imply distinct epigenetic regulatory functions for 5hmC, 5fC, and 5caC (Spruijt et al., 2013; Wu et al., 2014). Epigenetic changes are essential for normal development and frequently perturbed in cancer (Dawson, 2017). In particular peri-implantation development is accompanied by a dramatic epigenetic reprogramming entailing the global resetting of DNA modifications (Ficz et al., 2013; Habibi et al., 2013; Pfaffeneder et al., 2014).

Whereas the enzymes catalyzing the subsequent steps of DNA modification have been identified, still very little is known about their regulation and specific contributions during development. In particular the role of each of the three different TET proteins in the stepwise oxidation of 5mC at defined stages of cellular differentiation remains to be elucidated. Clearly, the observed stable cellular levels of oxidized cytosine derivatives and their distinct genome-wide distributions (Wu et al., 2014) seem to require dedicated regulatory mechanisms for each oxidation step. Currently available biochemical data do not conclusively resolve whether TET proteins oxidize 5mC in a processive manner or in a rather distributive mode with independent steps (Crawford et al., 2016; Tamanaha et al., 2016; Xu et al., 2014). Interestingly, the three TET proteins differ in their large, unstructured N-terminal domains possibly enabling divergent contributions to stage and cell type specific DNA modification (Iyer et al., 2009).

To systematically study the specific contribution of the three TET proteins, we used a well defined cellular differentiation system. The transition of cultured naive, pluripotent embryonic stem cells (ESCs) to primed-pluripotent Epiblast-like cells (EpiLCs) closely recapitulates peri-implantation development (Hayashi et al., 2011) with its dramatic epigenetic changes rendering it an ideal model system to study basic principles governing cytosine oxidation.

**Figure 1:**
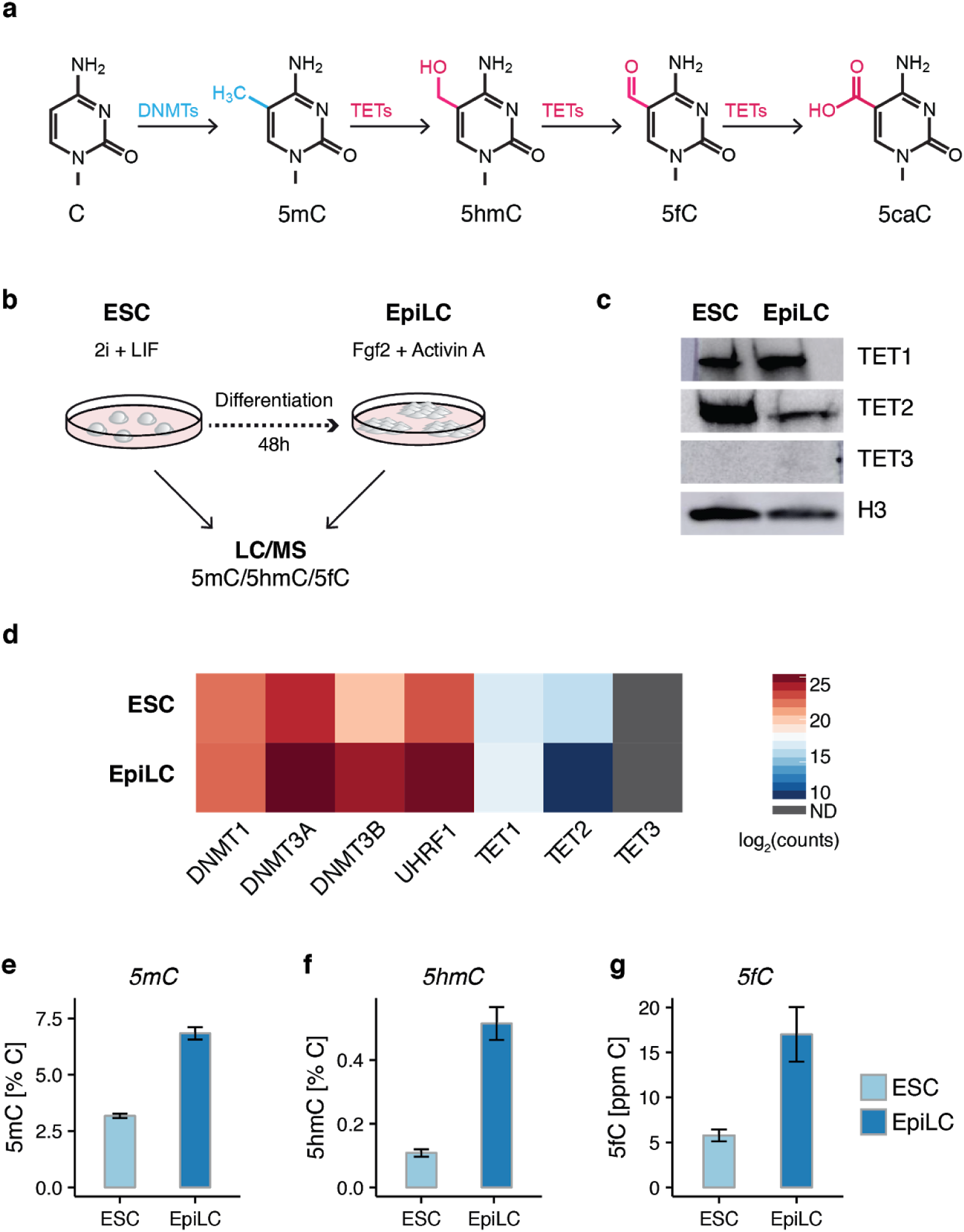
Quantification of cytosine derivatives and their respective enzymes in the transition from naive to primed pluripotency. (a) Cytosine modifications depicted with the enzymes responsible for their generation. (b) Schematic overview of experimental design. Samples for LC-MS/MS were collected at naive (ESC) and primed (EpiLC) pluripotency stages. (c) Western blot analysis of TET1, TET2, TET3 protein levels in wt ESCs and EpiLCs with histone H3 as loading control. (d) Heatmap depicting protein abundance of DNA modifying enzymes in ESCs and EpiLCs as determined by LC-MS/MS with protein abundance given as log2 peptide ratio. N.D.:no peptides detected. (e-g) Mean values ± standard deviation of DNA modification levels depicted as percentages of genomic cytosines in mESCs (*n* = 6) and mEpiLCs (*n* = 12).

To assess changes in DNA modifications during the transition of ESCs to EpiLCs, we quantified modified cytosines in genomic DNA with tandem liquid-chromatography mass spectrometry (LC-MS/MS) (Figure 1b). Global DNA methylation increased over the course of differentiation with 5mC levels in naive ESCs and EpiLCs closely resembling those of their respective *in vivo* counterparts, the E3.5 inner cell mass and E6.5 epiblast (Ficz et al., 2013) (Figure 1e; Supplementary Table 1). The precise quantification of cytosine derivatives demonstrated that along with 5mC also the levels of 5hmC and 5fC increased from ESCs to EpiLCs (Figure 1f-g; Supplementary Table 1). Interestingly, while 5mC levels doubled, 5hmC and 5fC even displayed a five- and threefold increase, respectively. This overproportional increase of 5hmC and 5fC suggests that the modification of cytosines occurs in successive steps subjected to independent regulation during exit form naive pluripotency.

In search of possible mechanisms, we examined how the six enzymes, known to be responsible for DNA modification in mammals, may contribute to these uncoupled levels of cytosine derivatives. Coinciding with the global wave of DNA methylation, mass spectrometry (MS)-based quantitative proteomics showed increased levels of the *de novo* DNA methyltransferases DNMT3A and DNMT3B during differentiation (Figure 1d), which is consistent with changes observed during peri-implantation development (Borgel et al., 2010). Whereas protein levels of the ubiquitous maintenance DNA methyltransferase DNMT1 remained constant, its essential regulator and cofactor UHRF1 increased and might thereby contribute to and maintain DNA methylation (Figure 1d). Interestingly, despite even larger gains in oxidized cytosine levels, we did not detect corresponding increases in TET protein levels during the transition from naive ESCs to EpiLCs. On the contrary, while TET1 levels remained relatively constant we measured a fortyfold reduction in TET2 peptides in EpiLCs and TET3 was not detectable in both cell types (Figure 1d). These changes in TET protein levels were directly confirmed by independent Western blot analyses (Figure 1c) and are consistent with mRNA levels determined by qPCR (Supplementary Figure 1a).

As these TET expression data cannot explain the observed changes in global DNA modifications and even show a negative correlation with 5hmC and 5fC levels, we considered changes in the Base Excision Repair (BER) pathway. Thymine DNA glycosylase (TDG) can excise genomic 5fC and 5caC that are then replaced by unmodified cytosine via BER and could thus change the abundance of modified cytosines in genomic DNA (Kohli and Zhang, 2013). However, our proteomics data from ESCs and EpiLCs indicated that levels of the BER pathway proteins (LIG3, PNKP, APEX1 and PARP1) remained unchanged while TDG levels even increased over differentiation (Supplementary Figure 1b). These data argue against reduced removal of oxidized cytosine derivatives by the TDG/BER pathway and, thus, only leave TET-mediated oxidation as a possible explanation for the observed increase in 5hmC and 5fC levels during the naive to primed pluripotency transition.

To dissect and identify the specific contributions of TET proteins at the transition from naive to primed pluripotency, we used CRISPR/Cas mediated mutagenesis to generate *Tet1* and *Tet2* single knockout (KO) and *Tet1/Tet2* double KO (DKO) ESC lines (Supplementary Figure 2a-b) and confirmed loss of TET1 and TET2 by Western blot analyses (Supplementary Figure 2c-d). Using two independent clones for each genotype, we quantified the levels of 5mC, 5hmC and 5fC in ESCs and EpiLCs by LC-MS/MS. Elimination of either TET1 or TET2, or both TET1 and TET2 resulted in modest yet significant increases in DNA methylation in both naive ESCs and primed EpiLCs (Figure 2a; Supplementary Table 1 and 2). The relatively minor 5mC gains in *Tet1* KO ESCs is consistent with a report that the bulk of the naive genome is kept hypomethylated via the inhibition of maintenance methylation, independent of TET proteins (von Meyenn et al., 2016).

**Figure 2.**
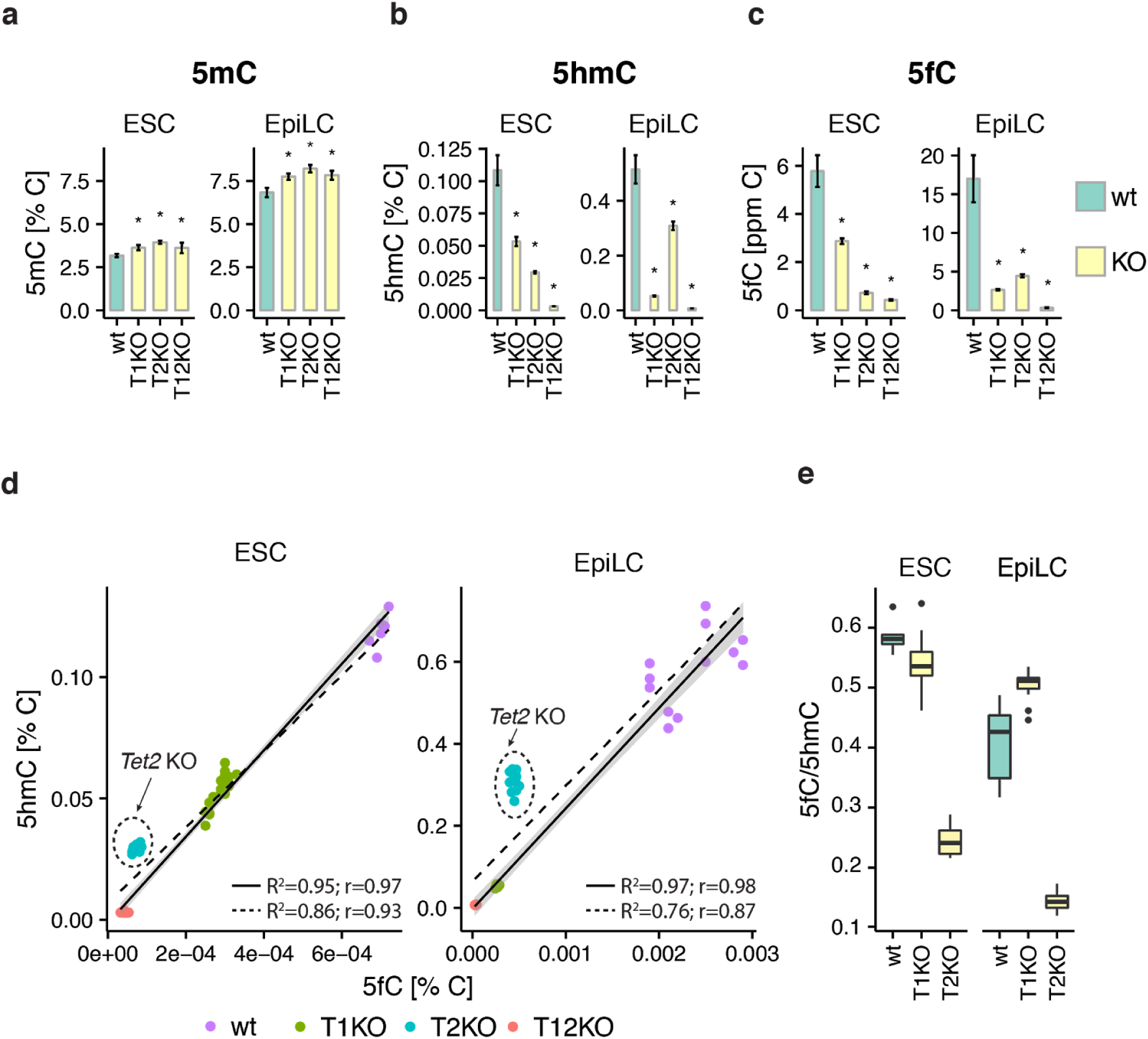
Quantification of cytosine modifications in *Tet1* and *Tet2* knockout ESCs and EpiLCs. (a-c) DNA modification levels as percentage of total cytosines in wt (*n* = 6 (ESCs); n = 12 (EpiLCs)), *Tet1* KO (T1KO; *n* = 22), *Tet2* KO (T2KO; *n* = 12), and *Tet1/Tet2* DKO (T12KO; *n* = 22) ESCs and EpiLCs. Depicted are mean values ± standard deviation; *P< 0.005 to wt as determined using a one-way ANOVA followed by a post-hoc Tukey HSD test. (d) Correlations between 5hmC and 5fC levels in wt and *Tet* KO ESCs and EpiLCs. The dashed regression line was generated using the full data set, the solid regression line was generated by excluding *Tet2* KO data. Depicted are values from the individual replicates presented in Figure 2a. R^2^: coefficient of determination; r: Pearson correlation coefficient. (e) Box plots of the ratio of 5fC to 5hmC in ESCs and EpiLCs in wt, *Tet1* KO and *Tet2* KO cell lines. The median is represented by the central bold line. The lower and upper hinges correspond to the first and third quartiles (the 25th and 75th percentiles). The upper and lower whisker extend from the hinge to the largest and lowest value, respectively, no further than 1.5 * interquartile range (IQR). Unlike the *Tet1* KO, *Tet2* KO drastically affects the 5fC/5hmC ratio.

Double *Tet1/Tet2* KO resulted in the loss of practically all oxidized cytosine derivatives with levels near the detection limit in ESCs and EpiLCs (Supplementary Table 2). Together with our expression data (Figure 1c-d; Supplementary Figure 1a) these results argue for a major role of TET1 and TET2 in 5mC oxidation during naive pluripotency exit with little to no contribution from TET3.

Analysis of the individual *Tet* KOs revealed stark, stage-specific differences in each enzyme’s functional contribution to the consecutive steps of cytosine oxidation. Genomic 5hmC levels were significantly decreased in *Tet1* KO (50% of wt 5hmC) as well as *Tet2* KO (30% of wt 5hmC) naive ESCs indicating that despite high expression levels both, TET1 and TET2, are not redundant (Figure 2b; Supplementary Table 2). The more dramatic 5hmC depletion in *Tet2* KO ESCs argues the majority of 5mC to 5hmC conversion in naive pluripotency to require TET2. Strikingly, *Tet2* KO 5hmC levels substantially increased upon exit from pluripotency and recovered from ∼0.03% (30% of wt ESC 5hmC) to ∼0.3% of genomic cytosines (60% of wt EpiLC 5hmC). Thus, TET1 largely appears to drive the differentiation-dependent acquisition of 5hmC (Figure 2b; Supplementary Table 1 and 2). Supporting this notion was the finding that *Tet1* KOs fail to acquire 5hmC upon exit from naive pluripotency, with 5hmC levels remaining essentially unchanged between naive and primed pluripotency (∼0.05% in ESCs and ∼0.06% in EpiLCs *Tet1* KO versus ∼0.5% of genomic cytosines in wt EpiLCs) (Figure 2b and Supplementary Table 1).

Despite the increased predominance of TET1 in 5hmC production during differentiation (Figure 2b) coincides with a massive downregulation of TET2 (Figure 1c-d), enzyme levels alone fail to provide mechanistic insights into the regulation of 5hmC formation in naive ESCs. Although naive ESCs express TET1 at levels similar to EpiLCs (Figure 1c-d) and possess far less 5hmC than in the primed state (∼0.1% versus ∼0.5% of genomic cytosines), TET1 in the absence of TET2 (in *Tet2* KO) is only able to sustain 30% of 5hmC in naive ESCs (Figure 2b and Supplementary Table 1). In other words, equal amounts of TET1 sustain ten-times less 5hmC in ESCs versus EpiLCs (∼0.03% versus ∼0.3% of genomic cytosines). Taken together, TET1 and TET2 possess distinct, stage-specific roles in the oxidation of 5mC, in which the responsibility of 5hmC formation passes from TET2 to TET1 upon differentiation.

To investigate whether similar stage-dependent preferences apply for the subsequent oxidation step, i.e. the conversion of 5hmC to 5fC, we compared 5fC levels in ESCs and EpiLCs. Analysis of 5fC levels in KO lines revealed an unexpected, prominent role of TET2 in ESCs and even EpiLCs. In naive ESCs, *Tet2* KO caused a loss of ∼87% in 5fC levels, almost reaching background levels of the *Tet1/Tet2* DKO, whereas only 50% of 5fC was lost in *Tet1* KO ESCs (Figure 2c). The reduction of 5fC in *Tet1* KO ESCs was proportional to the loss of its precursor 5hmC (Figure 2d-e). In striking contrast, the dramatic reduction of 5fC in naive *Tet2* KO ESCs did not correlate with a decrease in 5hmC (Figure 2d-e). Thus, TET2 is almost exclusively responsible for global cytosine oxidation in naive pluripotency and cannot be compensated for by TET1 in naive ESCs.

In EpiLCs, 5fC levels dropped to ∼18% and ∼26% of their wt levels in *Tet1/Tet2* KO cells, respectively (Figure 2c). Remarkably, the similarity of 5fC levels in both, *Tet1* and *Tet2* KO EpiLCs stands in stark contrast to their 5hmC levels (Figure 2b). Similar to naive ESCs, the major reduction of 5fC in *Tet1* KO EpiLCs is accompanied by a massive decrease in 5hmC. However, depletion of TET2 in EpiLCs led to a dramatically disproportionate decrease in 5fC compared to 5hmC (Figure 2d-e). This significant global depletion of 5fC resulting from TET2 loss in EpiLCs is particularly striking considering that TET2 is only barely expressed at this particular stage (Figure 1c-d, Supplementary Figure 1a).

As 5fC can be excised from DNA by the BER pathway, we investigated whether the decrease in 5fC in *Tet2* KOs might be an indirect consequence resulting from upregulation of DNA repair enzymes. We assessed the transcript levels of *Apex1, Parp1, Lig3, Pnkp* or *Tdg* at both time points using qPCR (Supplementary Figure 2e). Neither the depletion of TET2 nor TET1 significantly affected the expression of these genes in ESCs or EpiLCs. Therefore, the disproportionate decrease of 5fC in *Tet2* KOs appears to be a direct effect of TET2 loss. In conclusion, the disproportionate loss of 5fC in both stages, naive and primed, reveals a previously unappreciated prominence of TET2 in the formation of 5fC in pluripotent stem cells.

In summary, the systematic quantification of cytosine derivatives and their respective enzymes in this defined cellular differentiation system lead to a number of unexpected findings (Figure 3). Whereas the increase of 5mC during naive pluripotency exit correlated well with the growing abundance of the *de novo* DNA methyltransferases, DNMT3A and DNMT3B, the rising levels of oxidized cytosine derivatives, 5hmC and 5fC, were accompanied by stable TET1 and diminishing TET2 levels. In these cells TET3 seems to play little to no role, given its undetectable expression and the practically complete loss of genomic 5hmC and 5fC in cells lacking TET1 and TET2.

**Figure 3.**
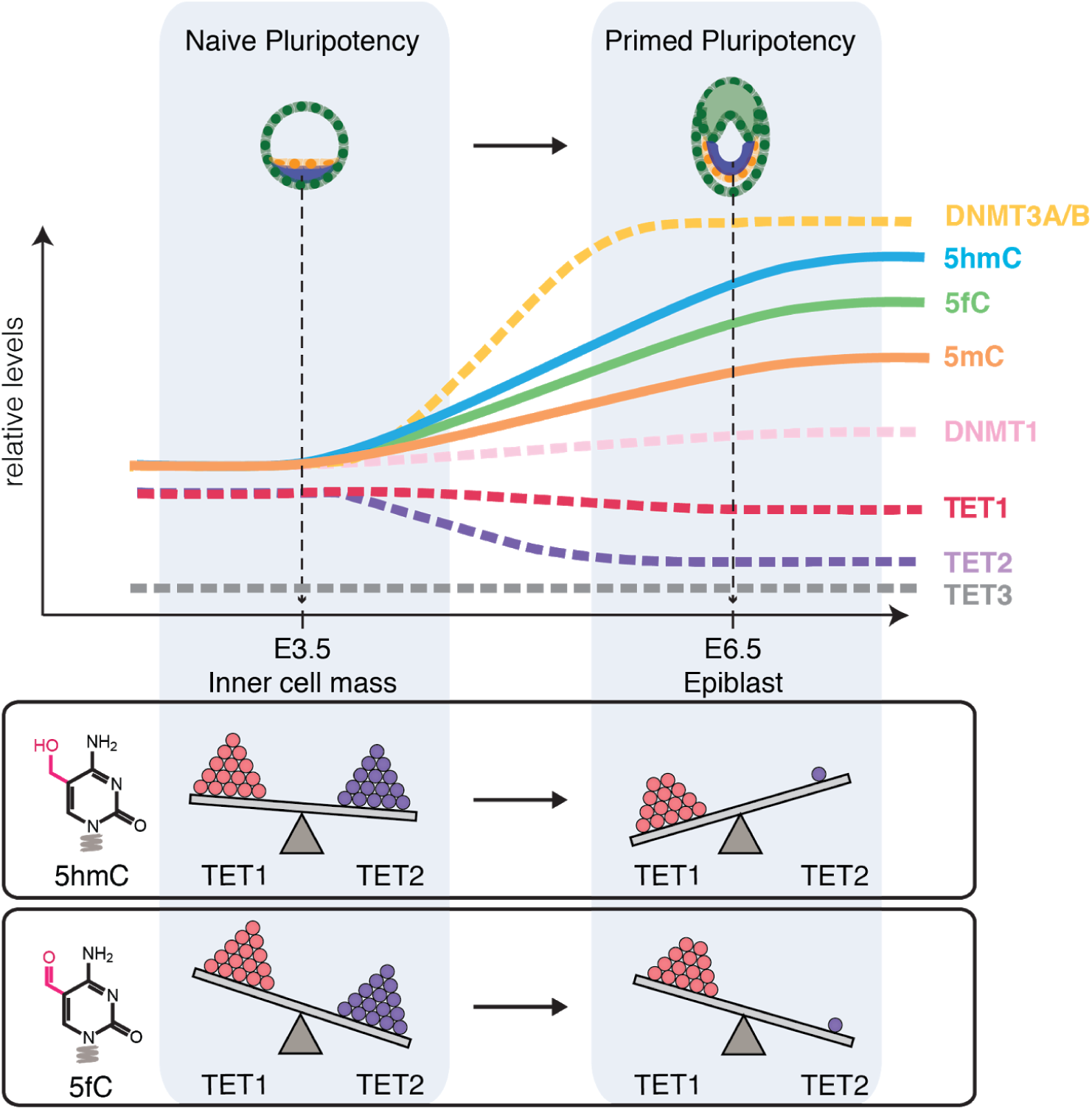
Epigenetic changes and distinct contributions of different DNA modifying enzymes during the transition from naive to primed pluripotency. Graphical summary depicting changes in cellular levels of cytosine modifications and their respective DNA modifying enzymes in the transition from naive to primed pluripotency. The relative contributions of TET1 and TET2 to the generation of 5hmC and 5fC as estimated from observations in *Tet* KO ESCs and EpiLCs are illustrated at the bottom; the number of spheres and tilt of the balance represent protein abundance and contribution to the cytosine derivative, respectively. TET1 gains importance in the oxidation of 5mC to 5hmC during differentiation as TET2 abundance decreases. Most remarkably, despite drastic downregulation TET2 remains critical for the formation of 5fC in primed pluripotency.

Our analysis of global cytosine modification levels in *Tet1* and *Tet2* KO ESCs and EpiLCs revealed both enzymes to have profound stage-specific contributions to cytosine oxidation which cannot be fully compensated by the other enzyme. In ESCs, the oxidation of 5mC to 5hmC relies primarily on TET2, whereas the global increase in 5hmC during differentiation is almost exclusively catalyzed by TET1.

The previously observed silencing had argued against any role of TET2 in peri-implantation development (Sohni et al. 2015; Khodeiry et al. 2017). We also observed downregulation of *Tet2* but still detected TET2 protein by mass spectrometry and Western blot analysis, and in fact our KO data identified a rather distinct role of TET2 at this stage. Remarkably, *Tet2* KO ESCs and EpiLCs show an unexpected loss of 5fC arguing that TET2 governs the formation of 5fC in ESCs and the pronounced accumulation of 5fC during the naive to primed transition. Our KO data clearly demonstrate that the residual amounts of TET2 proteins in EpiLCs have a prominent role in the oxidation of 5hmC to 5fC, which cannot be compensated by the much more abundant TET1.

Detailed analysis of the different KO lines also showed that 5hmC and 5fC levels respond quite independently. This apparent uncoupling of oxidized cytosine derivative levels together with the observed different roles of TET1 and TET2 in 5mC and 5hmC oxidation strongly argue against processive oxidation and favor DNA modification in three consecutive and independently regulated steps from 5mC to 5hmC to 5fC and finally 5caC. A similar division between TET1 and TET2 activities has been described for SALL4A-bound enhancers (Xiong et al. 2016). Our data suggest that the distinct roles of TET1 and TET2 in the consecutive steps of oxidation and their divergent regulation shape the genome-wide patterns of oxidized cytosine residues. Thus, modulation of the catalytic activity of three TET enzymes by differential isoform expression, posttranslational modifications, interacting factors, and site-specific recruitment could constitute an additional layer of epigenetic regulation. Interestingly, 5fC was recently revealed to possess novel characteristics such as the ability to distort the DNA double helix (Raiber et al., 2015) and directly mediate DNA-protein crosslinks (Ji et al., 2017; Li et al., 2017), with potentially far reaching consequences on transcriptional regulation and chromatin remodeling (Raiber et al., 2017). In this context, our observation that 5fC formation is largely TET2-dependent might also have novel implications for the understanding of *Tet2* mutations in cancerogenesis.

## Contributions

C.B.M. conceived and designed the study, performed experiments, analyzed the data, and wrote the paper. S.B. conceived, designed, and supervised the study, analyzed the data, and wrote the paper. H.L. designed and supervised the study and wrote the paper. F.R.T., E.P., and T.C. performed UHPLC-MS/MS measurements and evaluated the data. E.M.E. and M.S. generated and characterized the cell lines. M.M. performed the LC-MS/MS proteomics and analyzed the proteomics data. M.D.B. helped characterize the cell lines. P.S. and J.R. performed validation experiments. All authors read, discussed, and approved the manuscript.

## Acknowledgments

We thank Dr. Feng Zhang for providing the pSpCas9(BB)-2A-Puro (PX459) V2.0 (Addgene plasmid # 62988). We thank W. Qin, V.B. Ozan, and T. Tigney for help with experiments. This work was supported by the Deutsche Forschungsgemeinschaft (DFG) with grants SFB1064/A17 to H.L. and SFB1064/A22 to S.B.

M.B., M.M. and J.R. are fellows of the IMPRS-LS PhD program and P.S. of the Integrated Research Training Group (IRTG) of the SFB1064. F.R.T. thanks the Boehringer Ingelheim Fonds for a PhD fellowship.

## Methods

### Cell culture

Naive J1 mESCs were cultured and differentiated into EpiLCs as described previously (Hayashi and Saitou, 2013; Mulholland et al., 2015). In brief, for both naive ESCs and EpiLCs defined media was used, consisting of: N2B27 (50% neurobasal medium (Life Technologies), 50% DMEM/F12 (Life Technologies)), 2 mM L-glutamine (Life Technologies), 0.1 mM β-mercaptoethanol (Life Technologies), N2 supplement (Life Technologies), B27 serum-free supplement (Life Technologies), and 100 U/mL penicillin, 100 μg/mL streptomycin (PAA Laboratories GmbH). Naive ESCs were maintained on flasks treated with 0.2% gelatin in defined media containing 2i (1 μM PD032591 and 3 μM CHIR99021 (Axon Medchem, Netherlands)), 1000 U/mL recombinant leukemia inhibitory factor (LIF, Millipore), and 0.3% BSA (Gibco) for at least three passages before commencing differentiation.

For CRISPR-assisted cell line generation mESCs were maintained on 0.2% gelatin-coated dishes in Dulbecco’s modified Eagle’s medium (Sigma) supplemented with 16% fetal bovine serum (FBS, Biochrom), 0.1 mM ß-mercaptoethanol (Invitrogen), 2 mM L-glutamine, 1× MEM Non-essential amino acids, 100 U/mL penicillin, 100 μg/mL streptomycin (PAA Laboratories GmbH), homemade recombinant LIF tested for efficient self-renewal maintenance, and 2i (1 μM PD032591 and 3 μM CHIR99021 (Axon Medchem, Netherlands)).

To differentiate naive ESCs into epiblast-like cells, cells were plated on flasks treated with Geltrex (Life Technologies) diluted 1:100 in DMEM/F12 (Life Technologies) in defined medium containing 10 ng/mL Fgf2 (R&D Systems), 20 ng/mL Activin A (R&D Systems) and 0.1× Knockout Serum Replacement (KSR) (Life Technologies). Media was changed after 24 h and EpiLCs were harvested after 48 h.

Cells were regularly tested for Mycoplasma contamination by PCR.

### CRISPR/Cas-mediated gene knockout and Western blot

For the generation of *Tet1* and *Tet2* knockouts, *Tet1* and *Tet2*-specific gRNAs (Supplementary Table 3) were cloned into puromycin-selectable vector expressing both SpCas9 and gRNA (px459: F. Zhang Lab). mESCs were transfected with Cas9-gRNA vector using Lipofectamine 3000 (Invitrogen) according to manufacturer’s protocol. Two days after transfection, J1 mES cells were plated at clonal density in ESC media supplemented with 1 µg/mL puromycin (Gibco). Selection media was removed after 48 h, replaced with normal ESC media, and colonies were allowed to grow for an additional 4-5 days. Single ESC colonies were transferred into 96-well plates and the plates were duplicated after 2 days. Enrichment for mutated clones was accomplished by amplifying the CRISPR/Cas targeted region via PCR (oligonucleotides in Supplementary Table 3) and performing restriction-fragment length polymorphism (RFLP) analysis (Wang et al. 2013) with SacI or EcoRV (FastDigest; Thermo Scientific) for *Tet1* or *Tet2*, respectively (see also Supplementary Figure 2a). Cell lysis in 96-well plates, PCR on lysates, and restriction digest were performed as previously described (Mulholland et al., 2015).

Clones harboring biallelic mutations were then assessed for loss of TET1 or TET2 via Western blot. Western blots for both *Tet* KOs were performed as described previously (Mulholland et al., 2015) using monoclonal antibodies (rat anti-TET1 5D6, rat anti-TET2 9F7, and rat anti-TET3 23B9) (Bauer et al., 2015) and polyclonal rabbit anti-H3 (ab1791, Abcam) as loading control. Blots were probed with secondary antibodies anti-rat (112-035-068, Jackson ImmunoResearch) and anti-rabbit (170–6515, Bio-Rad) conjugated to horseradish peroxidase (HRP) and visualized using an ECL detection kit (Thermo Scientific Pierce).

### Quantitative real-time PCR (qRT-PCR) Analysis

Total RNA was isolated using the NucleoSpin Triprep Kit (Macherey-Nagel) according to the manufacturer’s instructions. cDNA synthesis was performed with the High-Capacity cDNA Reverse Transcription Kit (with RNase Inhibitor; Applied Biosystems) using 500 ng of total RNA as input. qRT-PCR assays with TaqMan probes or oligonucleotides listed in Supplementary Table 3 were performed in 10 µL reactions with 5 ng of cDNA used as input. For TaqMan and SYBR green detection, TaqMan Universal Mastermix (Applied Biosystems) and FastStart Universal SYBR Green Master Mix (Roche) were used, respectively. The reactions were run on a LightCycler480 (Roche).

### LC-MS/MS analysis of DNA samples

#### DNA digestion

Isolation of genomic DNA was performed according to earlier published work (Pfaffeneder et al., 2014). 1.0–5 μg of genomic DNA in 35 μL H_2_O were digested as follows: 1) An aqueous solution (7.5 μL) of 480 μM ZnSO_4_, containing 18.4 U nuclease S1 (Aspergillus oryzae, Sigma-Aldrich), 5 U Antarctic phosphatase (New England BioLabs) and labeled internal standards were added ([^15^N_2_]-cadC 0.04301 pmol, [^15^N_2_,D_2_]-hmdC 7.7 pmol, [D_3_]-mdC 51.0 pmol, [^15^N_5_]-8-oxo-dG 0.109 pmol, [^15^N_2_]-fdC 0.04557 pmol) and the mixture was incubated at 37 °C for 3 h. After addition of 7.5 μl of a 520 μM [Na]_2_-EDTA solution, containing 0.2 U snake venom phosphodiesterase I (Crotalus adamanteus, USB corporation), the sample was incubated for 3 h at 37 °C and then stored at ‒20 °C. Prior to LC/MS/MS analysis, samples were filtered by using an AcroPrep Advance 96 filter plate 0.2 μm Supor (Pall Life Sciences).

#### LC-MS/MS analysis

Quantitative UHPLC-MS/MS analysis of digested DNA samples was performed using an Agilent 1290 UHPLC system equipped with a UV detector and an Agilent 6490 triple quadrupole mass spectrometer. Natural nucleosides were quantified with the stable isotope dilution technique. An improved method, based on earlier published work (Pfaffeneder et al., 2014; Wagner et al., 2015) was developed, which allowed the concurrent analysis of all nucleosides in one single analytical run. The source-dependent parameters were as follows: gas temperature 80 °C, gas flow 15 L/min (N_2_), nebulizer 30 psi, sheath gas heater 275 °C, sheath gas flow 15 L/min (N_2_), capillary voltage 2,500 V in the positive ion mode, capillary voltage ‒2,250 V in the negative ion mode and nozzle voltage 500 V. The fragmentor voltage was 380 V/ 250 V. Delta EMV was set to 500 V for the positive mode. Compound-dependent parameters are summarized in Supplementary table 1. Chromatography was performed by a Poroshell 120 SB-C8 column (Agilent, 2.7 μm, 2.1 mm × 150 mm) at 35 °C using a gradient of water and MeCN, each containing 0.0085% (v/v) formic acid, at a flow rate of 0.35 mL/min: 0 →4 min; 0 →3.5% (v/v) MeCN; 4 →6.9 min; 3.5 →5% MeCN; 6.9 →7.2 min; 5 →80% MeCN; 7.2 →10.5 min; 80% MeCN; 10.5 →11.3 min; 80 →0% MeCN; 11.3 →14 min; 0% MeCN. The effluent up to 1.5 min and after 9 min was diverted to waste by a Valco valve. The autosampler was cooled to 4 °C. The injection volume was amounted to 39 μL. Data were processed according to earlier published work (Pfaffeneder et al., 2014).

### MS-based quantitative proteomics

#### FASP-based protein digestion

10 µg proteome samples were digested with a modified FASP procedure (Wiśniewski et al., 2009). First, proteins in lysis buffer were reduced and alkylated by the addition of 40 mM of IAA for 45 min in darkness at room temperature (RT). One volume of 8 M Urea diluted with 0.1 M TRIS/HCl pH 8.2 and 10% isopropanol in 100 mM TEAB was then added to the protein sample. 30 kDa molecular weight cut-off filter devides (PALL, Port Washington, USA) were used and samples were washed twice with UA buffer and twice with 50 mM ammonium bicarbonate prior to digestion. Protein digestion was then performed by adding 1 µg Lys-C (Wako Chemicals) to the immobilized proteins on the filter and incubating for 2 h at RT, followed by 1 μg of trypsin (Promega) in 50 mM TEAB overnight incubation at RT. Peptides were recovered using an initial spin of 3,100 × g for 10 min followed by two centrifugations with 50 μL of 50 mM TEAB. Recovered peptides were stored at -20 °C.

#### Proteomics MS measurements

The LC‒MS/MS experiments were performed on the Ultimate 3000 nano-RSLC (Thermo Scientific) system coupled with the high resolution QExactive HF mass spectrometer (Thermo Scientific) (Thermo Scientific). Individual peptide fractions were reconstituted in 3% acetonitrile, 0.1% formic acid and loaded on the Acclaim PepMap RSLC, 75 μm × 25 cm, nanoViper, C18, 2 μm particle column for multistep gradient elution with the injection mode using the loading pump at 30 μL/min flow rate for 5 min. The gradient elution method at flow rate 300 nL/min was as follows: for 90 min gradient up from 5% to 25%, followed for 5 min gradient up to 40% in 0.1% FA. Top 10 multiply charged precursor isotopic clusters with m/z value larger than 300 or smaller than 1500 were selected for fragmentation with FT mass resolution of 60000 and isolated for HCD fragmentation within a mass window of 1.2 Da. Normalized collision energy was set to 27 an isolation window of 1.6 m/z and a dynamic exclusion of 30sec.

#### Processing proteomics data

The MS spectra were analyzed using Proteome Discoverer 1.4 (Thermo Fisher Scientific), and the proteins were identified using MASCOT search engine (Matrix Science) against the *Mus musculus* proteome of the Uniprot database. Searches were carried out based on tryptic specificity. The precursor and ms/ms tolerance were set on 10 ppm and and 20 mmu fragment mass tolerance, respectively. Carbamidomethylation of Cytosine was selected as a fixed modification, and as dynamic modifications, methionine oxidation and asparagine or glutamine deamidation were selected. The Perculator algorithm (Käll et al., 2007) was used to limit FDR rates to a q-value<0.01. Proteins were identified and quantified by at least 1 unique peptide. For subsequent analysis, protein grouping was enabled to consider only the master proteins. An estimated peptide false discovery rate (FDR) identification was set to 1% using a Mascot search engine. In addition, peptide assignments were re-imported into the Progenesis QI software and the abundances of all peptides allocated to each protein were summed up.

